# Viral silencing suppressor activity in plants modifies aphid antiviral immunity and fecundity

**DOI:** 10.1101/2025.05.13.653761

**Authors:** Stephanie E. Preising, Jennifer R. Wilson, Glenn M. Parker, Stacy L DeBlasio, Domenica Reeve, Joshua S Chappie, Michelle L Heck

## Abstract

Plants defend themselves from viral infection using RNA interference (RNAi), an evolutionarily conserved mechanism that degrades viral RNA through the production of small interfering RNAs. As a counter defense, plant viruses evolved suppressors of RNA silencing (VSRs) to inhibit plant RNAi machinery, aiding viral replication and transmission. P0, a VSR encoded by the potato leafroll virus (PLRV), family *Polerovirus*, suppresses RNAi by targeting the plant protein ARGONATE 1 for degradation through its F-box motif interaction with a Skp1 subunit of the family of E3 ubiquitin ligases. Our previous work shows PLRV P0 suppresses antiviral immunity in its aphid vector *Myzus persicae*, leading to an increase in aphid infection by the insect virus Myzus persicae densovirus (MpDNV). Here, we expand on these findings and show that the P0 protein also regulates aphid fecundity. Using a series of F-box mutants, we demonstrate that a functional P0 F-box motif is required for inhibition MpDNV antiviral immunity but not modulation of aphid fecundity. We further show that silencing suppressors from non-aphid-borne plant viruses that target other components of the plant’s RNAi machinery also modulate aphid fecundity but not MpDNV titer. Collectively, the results show that aphids have been favored by selection to modulate their anti-viral immunity and fecundity in response to changes in the plant RNAi pathways induced by plant viral infection. These data highlight the intricate co-evolution of plant viruses, their vectors, and host defenses. This knowledge may open new avenues for managing vector-borne plant diseases by targeting viral proteins to manipulate insect vectors.

## Introduction

To thwart RNA virus infection, plants and insects use RNA interference (RNAi), a conserved evolutionary defense mechanism for the targeted destruction of viral mRNA molecules. Dicer-like proteins, a family of ribonucleases that cleave viral double-stranded RNA (dsRNA) intermediates of viral replication, produce a population of small interfering RNAs (siRNAs) typically ranging in size from 21-24 nucleotides (Lindbo and Dougherty 2005). The siRNAs are loaded into RNA-induced silencing complexes (RISC) where they engage ARGONAUTE (AGO) proteins, providing the complimentary base pairing needed to direct the sequence-specific degradation of viral RNA (Ding and Voinnet 2007), halting translation of viral replication products. The anti-viral RNAi response in plants confers systemic immunity. Insects, including aphids, use a similar pathway but with limited systemic spread in the insect body and varying degrees of efficiency in different insect species (Sukhikh et al. 2024).

As a counter defense to RNAi, most viruses encode viral suppressors of RNA silencing (VSRs), that disrupt the host’s RNAi pathways (Ahn et al. 2011; Bortolamiol et al. 2007; Burgyán and Havelda 2011; González et al. 2010). In plants, VSRs encompass genetically diverse sequences with highly conserved functions in their targeting of the plant RNAi pathway components, consistent with convergent evolution of these counter-defense proteins. Some VSRs prevent the assembly of the effector RISC by targeting its components, notably AGO-1 (Bortolamiol et al. 2007). For example, the 2b protein of cucumber mosaic virus (CMV-2b, family *Bromoviridae*) interacts directly with the conserved AGO1 domains PAZ and PIWI, essential for loading small RNA (sRNA)s and retaining endonuclease activity, respectively, by inhibiting the cleavage activity of these enzymes (Duan et al. 2012; Zhang et al. 2006; Zhao et al. 2023). The tomato bushy stunt virus (TBSV, family *Tombusviridae*) p19 VSR inhibits the dsRNA to siRNA pathway by sequestering and eradicating the use of vsiRNA (virus-derived siRNAs) and by preventing loading onto the RISC complex (Ahn et al. 2011). Consequentially, vsiRNAs used as systemic signals in plant antiviral silencing do not reach distant and uninfected parts of the plant. The turnip crinkle virus (TCV, family *Tombusviridae*) p38 VSR inhibits the processing of siRNA through AGO1 binding, affecting the stability of the RNAi network, and ultimately inhibiting Dicer-like proteins (Iki et al. 2017).

The potato leafroll virus (PLRV, family *Solemoviridae*), encodes the VSR P0 in ORF 0 of its single-stranded, positive-sense RNA genome. P0 is an F-box protein that suppresses the plant’s antiviral defense by targeting AGO1 for degradation in a proteasome-independent manner by promoting the formation of ER-related bodies that deliver P0 and AGO1 to the vacuole, allowing PLRV proliferation (Bortolamiol et al. 2007). While the polerovirus P0 protein function is conserved across virus species, the amino acid sequences of P0 proteins from different polerovirus species are diverse, with an average amino acid sequence identity of 37.1% in the *Polerovirus* genus (Barrios Barón et al. 2021). The conserved regions in the F-box domain are critical for VSR function (Bortolamiol et al. 2007; Rashid et al. 2019). Single amino acid changes in the F-box alters the silencing suppressor function and virus-host specificity (Bortolamiol et al. 2007; Burgyán and Havelda 2011; Cascardo et al. 2015). The generation of PLRV P0 F-box mutants in an infectious clone targeting P0 amino acids 76-95, and P0 mutants with modifications to the C-terminal domain show that P0 is necessary for virion accumulation and systemic infection (Rashid et al. 2019).

PLRV is exclusively transmitted in a circulative, non-propagative manner by the green peach aphid, *Myzus persicae* (Hemiptera: *Aphididae*). Circulative, non-propagative transmission of PLRV involves aphid acquisition of virions from the phloem of infected plants during prolonged periods of phloem-feeding (Wilson et al. 2020). It is unknown whether aphids ingest P0 or other PLRV proteins during virus acquisition from systemically-infected plant tissue. However, expression of P0 in plants has been shown to impact interactions with aphid vectors (Patton et al. 2020). Transient plant expression of PLRV P0 increases the fecundity of the PLRV aphid vector, *Myzus persicae*, and impacts the aphid’s immune response to insect-infecting viruses (Patton et al. 2020; Pinheiro et al. 2019).

PLRV-viruliferous *M. persicae* aphids display a modified anti-viral immune response to Myzus persicae densovirus (MpDNV), an aphid-infecting DNA virus in the family *Parvoviridae*, transmitted among aphids through saliva, honeydew, and transovarially (Van Munster et al. 2003). PLRV-viruliferous aphids that acquire PLRV from infected plants display an unusual sRNA size profile for sRNA reads mapping to the MpDNV genome. Specifically, PLRV-viruliferous aphids produce an abundance of long sRNA species, averaging 35 nucleotides long, and a reduction in the number of 22-mers matching to the MpDNV genome (Pinheiro et al. 2019). This sRNA profile is unique to PLRV-viruliferous aphids that have acquired virus from infected plants. Aphids collected from healthy plants, PLRV-containing, or sucrose-only artificial diets, and plants infected with the non-persistent, aphid-borne virus Potato virus Y (PVY) produced the expected size of sRNAs matching to the MpDNV genome (Pinheiro et al. 2019). The MpDNV titer is higher in PLRV-viruliferous aphids as compared to non-viruliferous aphids, supporting the hypothesis that PLRV impairs the aphid’s antiviral RNAi immune signaling pathway. Further, expression of PLRV P0 alone in the absence of PLRV infection also increases MpDNV titer in aphids, suggesting the silencing suppression activity of PLRV P0 in plants is responsible for suppressing the aphid’s antiviral immune response against MpDNV (Pinheiro et al. 2019). Intriguingly, the effect of PLRV on the aphid’s immune system extends beyond DNA viruses but also to RNA viruses. We discovered that viruliferous *M. persicae* produce no sRNAs against Myzus persicae flavivirus (MpFV) (Larrea-Sarmiento et al. 2024). Thus, the PLRV-induced changes in sRNA production for DNA and RNA viruses in viruliferous aphids is distinct, with a size shift to larger sRNAs mapping to MpDNV and a complete inhibition of sRNA production mapping to MpFV.

Insect-infecting viruses also encode VSRs. Flock House Virus B2 silencing suppressor protein binds and sequesters siRNA intermediates, preventing the loading and binding by Ago-2 containing complexes (Singh et al. 2009). The 1A protein from cricket paralysis virus (CrPV) suppresses RNA silencing with its indirect interaction with AGO2 in *Drosophila melanogaster* (Nayak et al. 2010). In aphids, MpDNV may be targeted by the aphid’s siRNA and pi-RNA pathways (Sukhikh et al. 2024). Many components of the RNAi pathway are conserved in aphids with notable expansions in pasha, dicer-1, and AGO-1 gene families (Pea Aphid Genome Consortium 2010). These gene family expansions are present in multiple aphid species (Jaubert-Possamai et al. 2010) but appear to be specific to aphids among all metazoan species (Pea Aphid Genome Consortium 2010). The antiviral RNAi response occurs in multiple tissues in aphids, including the gut and the salivary glands. However, the overall RNAi response in aphids is weaker compared to *D. melanogaster* and other insects. While no evolutionary mechanisms have been proposed to explain phenotypic variation in RNAi among different insect species, it is intriguing to speculate that the RNAi pathway in hemipteran insects co-evolved with their plant hosts. This hypothesis highlights a possible role for VSRs to modulate antiviral immunity of both plant hosts and insect vectors.

In this study, we explore this hypothesis at the molecular and organismal levels. First, we tested whether the P0-dependent mechanism used by PLRV to modulate aphid anti-viral immunity depends on residues in the P0 F-box motif responsible for the silencing suppressor activity *in planta.* To evaluate whether the ability of plant viral SSRs to modulate aphid anti-viral defense was specific to PLRV P0, we also measured MpDNV titer in aphids after expression of other types of plant viral SSRs that target different proteins in the plant RNAi pathway. We serendipitously discovered that plant VSRs targeting different components of the plant RNAi pathway also impact aphid fecundity, suggesting that aphids have been favored by selection to rapidly respond to changes in the plant’s antiviral immune signaling pathways in very short time frames (within a single generation). Given that approximately 80% of plant viruses depend on insect vectors for plant-to-plant transmission (Hohn 2007), understanding the molecular tactics encoded in plant viral genomes that facilitate manipulation of insect vector phenotypes may provide new ways to manage the spread of insect vector-borne plant viruses (Blua et al. 1994; Chesnais et al. 2019; Heck and Brault 2018; Ingwell et al. 2012; Pinheiro et al. 2019).

## Results

### PLRV P0 F box mutants exhibit a range of silencing suppression activity in plants

Given that the F-box domain is crucial for P0 function in plants (Bortolamiol et al. 2007) and P0 expression in plants modulates aphid anti-viral immunity to MpDNV (Pinheiro et al. 2019), we tested whether the P0 F-box domain is required for VSR activity in aphids. The amino acid sequences of polerovirus P0 proteins are highly divergent, which has led to discrepancies in how the F-box motif is defined across different homologs (Pazhouhandeh et al. 2006) (Sun et al. 2020). To remove potential alignment biases and provide a more accurate framework for mutagenesis, we generated an AlphaFold (Jumper et al. 2021) model for PLRV P0 and compared it to the crystallized structure of human Skp2 (PDB: 1FQV), a prototypical F-box protein that functions as a substrate-recognition module in SCF ubiquitin-protein ligases (Gstaiger et al. 2001) (Supplemental Fig. S1). In Skp2, the F-box motif forms a compact helical bundle (Supplemental Fig. S1C and S1E). AlphaFold predicts a similar helical arrangement for the putative F-box region in PLRV P0 but with an additional β-hairpin inserted between the α1 and α2 helices (confidence >90%) (Supplemental Fig. S1D and S1E). Accounting for this insertion corrects prior ambiguities and yields an alignment where the F-box consensus motif can be clearly identified in each polerovirus P0 protein analyzed (Fig. 1 and Supplemental Fig. S1E).

**Figure 1.**
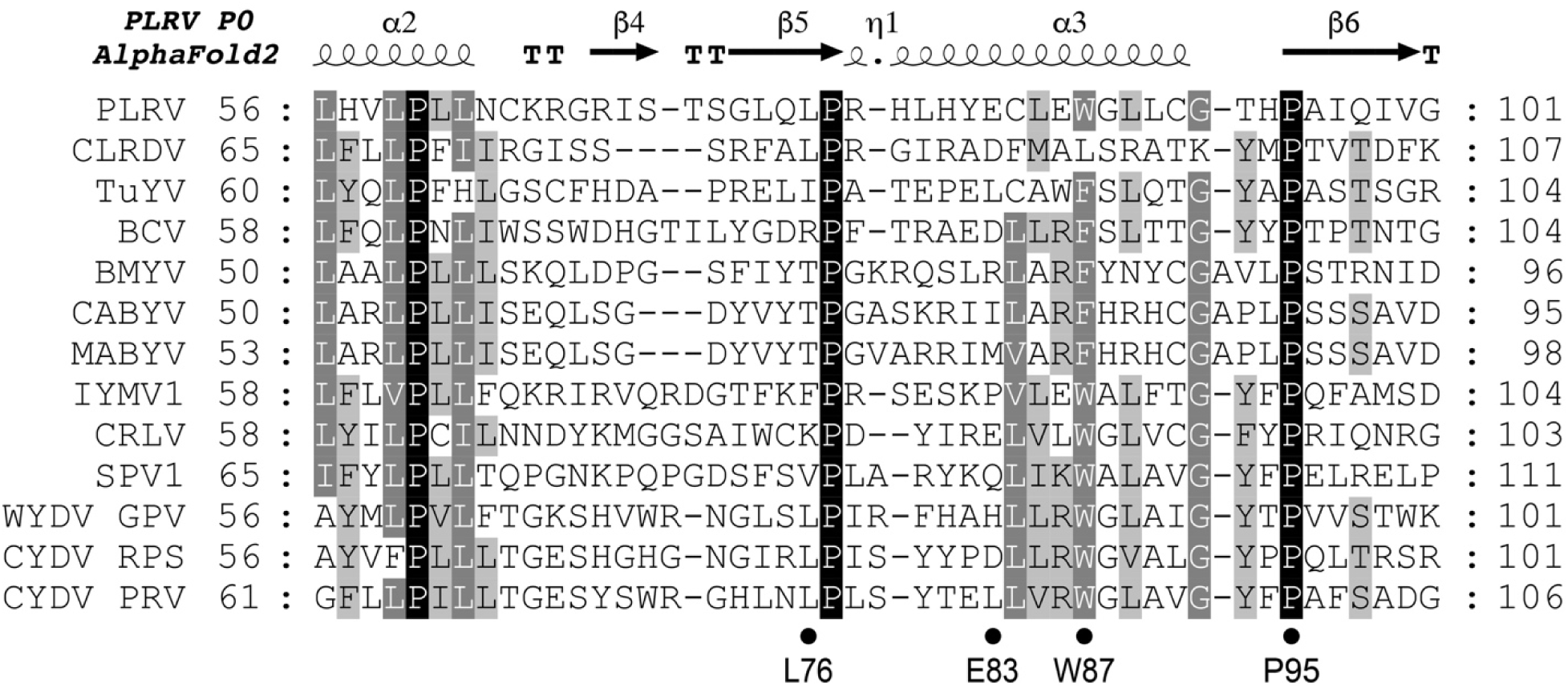
Polerovirus P0 amino acid sequence alignments show conserved and variable regions within the F-box domain. A sequence alignment of the F-box domain segment from representative polerovirus P0 proteins. The shading indicates sequence conservation across these homologs: white text on black background, 100% conserved; white text on dark gray background, 80% conserved; black text on light gray background, 60% conserved(Jumper et al. 2021). Filled black circles below the alignment designate PLRV P0 point mutations tested for silencing suppressor activity. Abbreviations are as follows with accompanying KEGG IDs, PLRV, potato leafroll virus (vg:1493890); CLRDV, cotton leafroll dwarf virus (vg:9777809); TuYV, turnip yellows virus (vg:940485); BCV, beet chlorosis virus (vg:921082); BMYV, beet mild yellowing virus (vg:935282); CABYV, cucurbit aphid-borne yellows virus (vg:940450); MABYV, melon aphid-borne yellows virus (vg:6369692); IYMV1, ixeridium yellow mottle virus 1 (vg:27111913); CRLV, carrot red leaf virus (vg:3021800); SPV-1, strawberry polerovirus 1 (vg:22276106); WYDV GPV, wheat yellow dwarf virus GPV (vg:10220414); CYDV RPS, cereal yellow dwarf virus RPS (vg:1489892); CYDV RPV, cereal yellow dwarf virus RPV (vg:1478314) (Kanehisa et al. 2017).

To test the if the P0 F-box motif has an impact on silencing suppressor activity in aphids, we cloned and expressed P0 mutants into a binary vector for transient, in planta expression of new (E83A and P95A) and previously published (W87A, L76A and L76R) (Rashid et al. 2019; Zhuo et al. 2014) single amino acid substitution mutants. To assess the silencing suppressor activity of the P0 F-box mutants, we used transient expression assays to evaluate the impact of the mutations on F-box activity in plants. Using *Agrobacterium tumefaciens*-mediated DNA delivery, we transiently co-expressed the PLRV P0 wild type (wt) or P0 mutants with vectors expressing GFP and a double stranded DNA sequence matching to the GFP gene (dsGFP), in a 1:1:1 ratio. GFP expression in the leaves was photographed three days post infiltration (DPI) via a long-wavelength UV lamp (Fig. 2). The GFP intensity at the cellular level was captured and quantified using a confocal microscope (Supplemental Fig. S2). In these assays, we co-expressed a 1:1 mixture of *A. tumefaciens* cells containing the GFP and dsGFP plasmids, which resulted in GFP silencing. This combination was used as our negative control. Analysis of the negative control resulted in a near-total GFP silencing with an average fluorescence intensity of 0.385 relative fluorescence unites (RFU) (Fig. 2). An increase in GFP fluorescence greater than this background level indicates functional silencing suppression, as shown by the expression of P0wt (Supplemental Fig. S3) when mixed in the 1:1:1 ratio with dsGFP and GFP (average fluorescence intensity = 21.537*, P* < 0.001). The P0 mutant E83A, located on the surface of the P0 protein (Supplemental Fig. 1), results in significantly higher GFP fluorescence at four DPI as compared to the dsGFP/GFP control (average fluorescence intensity = 29.197, *P* < 0.001). In contrast, L76A, P95A, L76R, and W87A does not significantly increase GFP fluorescence over the dsGFP/GFP control, (average fluorescence intensities; 4.814, 5.23933, 0.830, 0.132 RFU, *P-*values; (0.724, 0.540, 1.000, 1.000) (Fig. S3), although some fluorescence is detected. These mutations, predicted to be buried within in the P0 structure (Supplemental Fig. 1), reduce the P0 silencing suppression activity (Table 1). Collectively, these experiments allowed us to determine which P0 mutants impaired (L76A, P95A, L76R and W87A) or retained (E83A) F-box function to advance to testing their activity in aphids.

**Figure 2.**
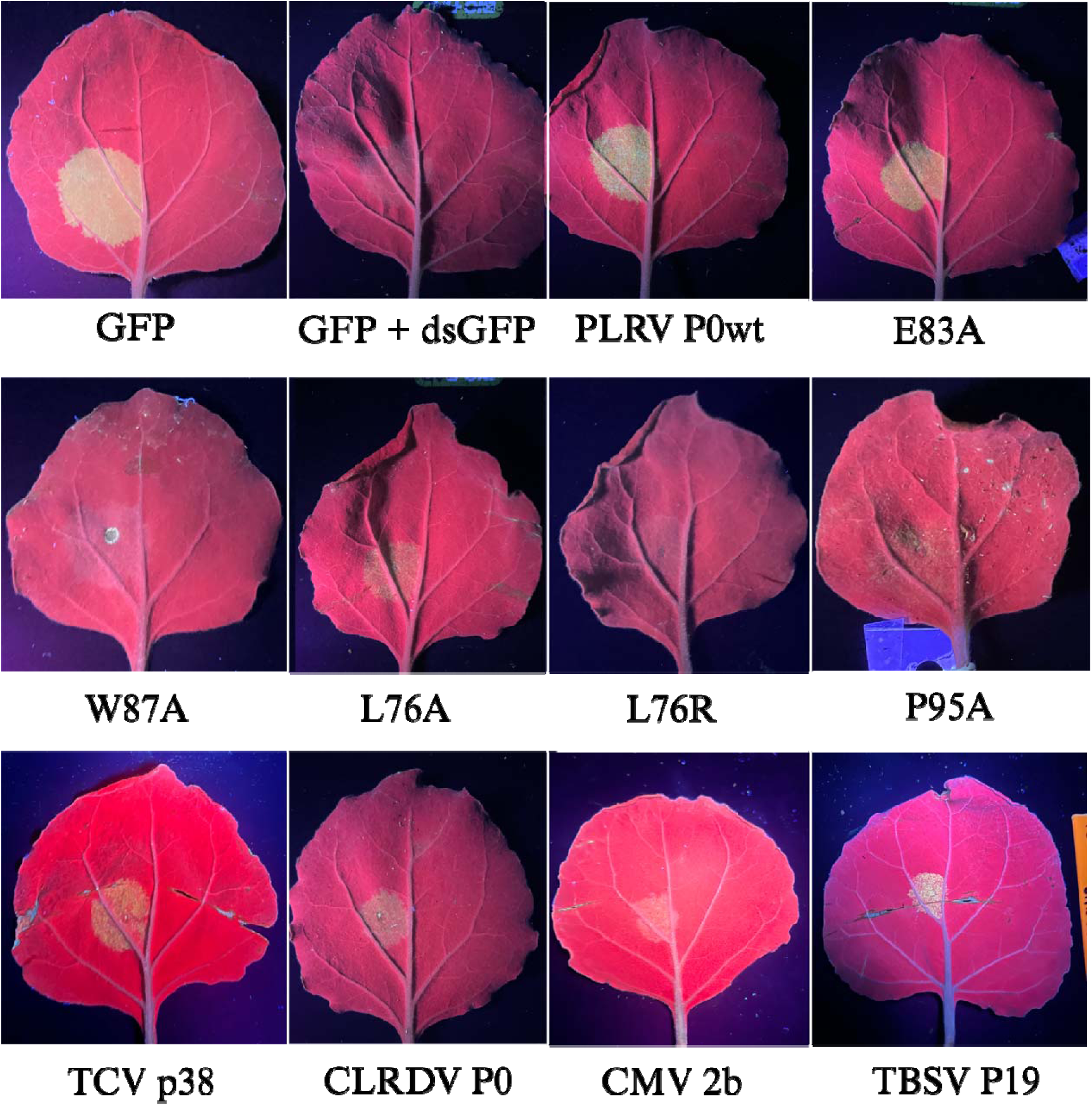
The silencing suppressor capability of PLRV P0 wt, PLRV P0 mutants, and a panel of VSRs transiently expressed in *Nicotiana benthamiana* leaves. The abaxial side of *N. benthamiana* leaves were co-inoculated with *Agrobacterium tumefaciens* vectors expressing the PLRV P0 wt construct, PLRV-P0 mutans, TCV 38p, CLRDV P0, TBSV P19, CMV 2b, together with *Agrobacterium* vectors expressing GFP and dsGFP, in a 1:1:1 ratio to evaluate the suppressor activity. For controls, *N. benthamiana* leaves were infiltrated with an *Agrobacterium* vector containing only GFP, and with GFP and dsGFP in a 1:1 ratio. Photographs were taken at three days post infiltration under a long-wavelength UV lamp.

**Table 1.** Average fluorescence intensity of GFP for each silencing suppressor protein as observed by confocal microscopy.

| Description | Treatment | Average<br>Fluorescence<br>intensity (RFU) | GFP expression<br>by confocal | *P*-value<sup>1</sup> |
| --- | --- | --- | --- | --- |
| Positive control | GFP | 47.333 | + | *P*<0.001 |
| Negative control | dsGFP/GFP | 0.385 | – |  |
| Silencing suppressor | GFP/dsGFP/PLRV P0 | 21.537 | + | *P*<0.001 |
|  | GFP/dsGFP/E83A | 29.197 | + | *P*<0.001 |
|  | GFP/dsGFP/L76R | 0.132 | – | 1.000 |
|  | GFP/dsGFP/P95A | 5.3933 | +<sup>2</sup> | 0.540 |
|  | GFP/dsGFP/W87A | 0.830 | – | 1.000 |
|  | GFP/dsGFP/L76A | 4.814 | +<sup>2</sup> | 0.724 |
|  | GFP/dsGFP/TVC p38 | 27.673 | + | *P*<0.001 |
|  | GFP/dsGFP/CLR DV P0 | 6.493 | + | 0.230 |
|  | GFP/dsGFP/CMV 2b | 12.788 | +<sup>2</sup> | *P*<0.001 |
|  | GFP/dsGFP/ TBSV P19 | 17.739 | + | *P*<0.001 |

1. *P*-values based on ANOVA Tukey's post hoc test as compared to the dsGFP/GFP control.
2. GFP visible by confocal microscopy with average fluorescence intensity not statistically different from dsGFP/GFP control.

### F-box motif activity is required for P0-mediated changes in MpDNV titer in *M. persicae*

We repeated the experiments described in Pinheiro *et al*. (2019), and showed aphids feeding on plant tissue transiently expressing PLRV P0 have a higher titer of MpDNV as compared to the GFP and un-infiltrated controls (Supplementary Fig. S4), which allowed us to proceed in testing the P0 mutants for effects on regulating MpDNV titer. We tested if P0 and P0 F-box mutant proteins alter the titer of MpDNV in aphids by caging aphids onto *N. benthamiana* leaves transiently expressing six different P0 treatments, a GFP control, and an un-infiltrated control as follows: (1) PLRV P0wt, (2) P0 E83A, (3) P0 P95A, (4) P0 L76A, (5) P0 L76R, (6) P0 W87A. After two days of exposure (equal to three DPI), the PLRV P0wt treatment resulted in aphid MpDNV titer that is significantly higher than P0 P95A, W87A, GFP, and the un-infiltrated control (Kruskal-Wallis test, *P* < 0.05, Fig. 3). There is no significant difference in MpDNV titer between aphids feeding on P0wt or E83A, the P0 mutant that does not interfere with silencing suppression activity in plants, or between L76A and L76R (Kruskal-Wallis test, *P* > 0.05, Fig. 3a). Aphids feeding on the mutants L76R and L76A have a lower average MpDNV titer as compared to P0 wt with higher variability and the difference in titer is not significant (Fig. 3; L76A *P* = 0.153, L76R *P* = 0.152). Collectively, these results support the hypothesis that the silencing suppressor activity of P0 in plants mediated by F-box activity is necessary to modulate MpDNV titer in aphids.

**Figure 3.**
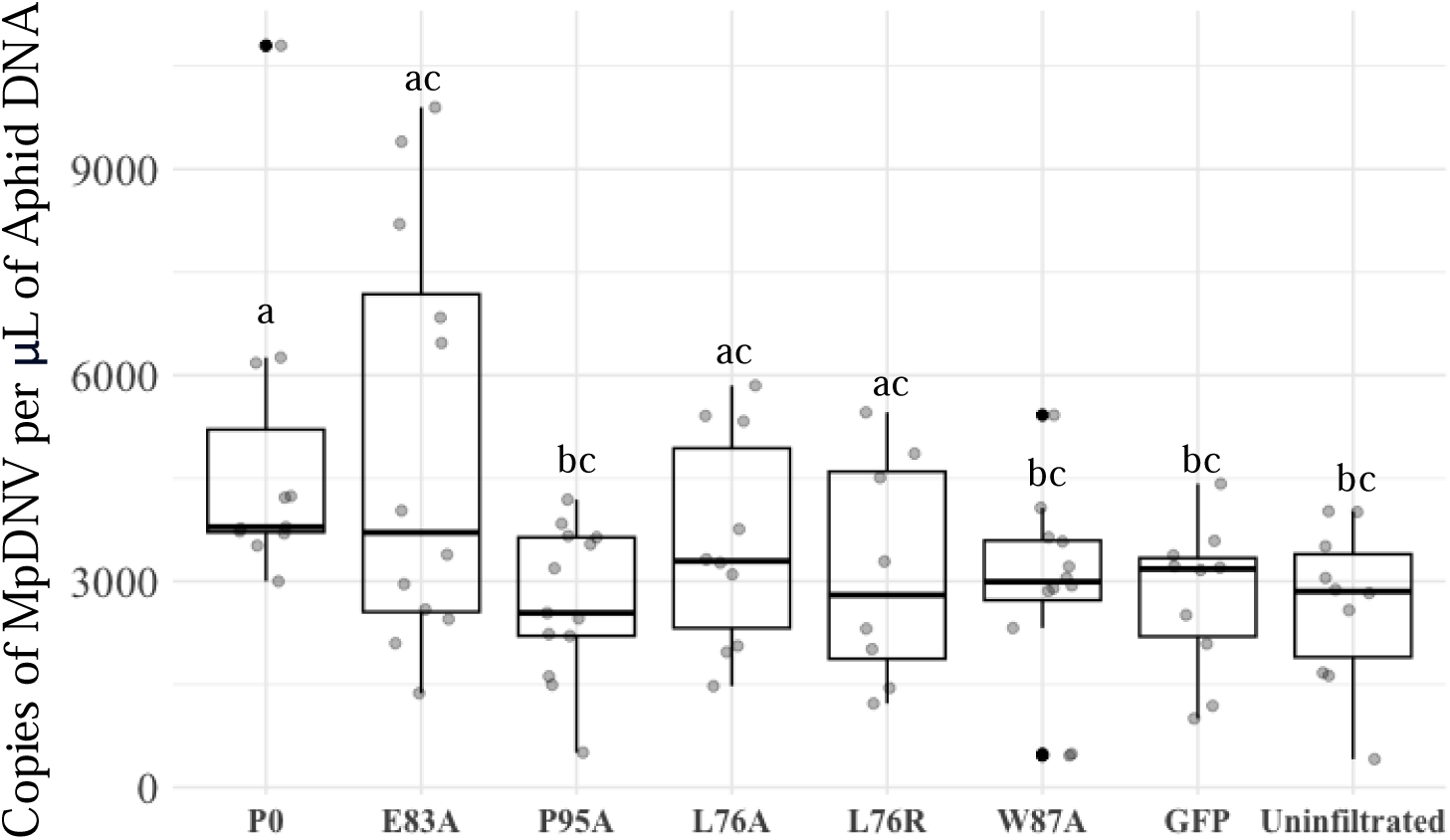
F-box motif activity modulates P0-mediated changes in MpDNV titer in *M. persicae*. The MpDNV titers of aphids on *N. benthamiana* plants transiently expressing PLR P0 wt and PLRV P0 mutants. Aphids were synchronized and clip caged on *N. benthamiana* transiently expressing P0 wt, P0 mutants, GFP, and an un-infiltrated control for two days. Aphid DNA was extracted and MpDNV titers were quantified via droplet digital PCR, with three pooled aphids in each replicate, and at least ten replicates per treatment over multiple trials. Aphid DNA was extracted and MpDNV titer was quantified via droplet digital PCR, three pooled aphids in each replicate, at least ten replicates per treatment. Letters indicate significantly different treatments via Kruskal-Wallis test.

### PLRV P0 regulates aphid fecundity independent of its silencing suppression function in plants

A preliminary experiment showed that *M. persicae* has a higher reproductive rate on PLRV-infected potato plants (not shown), which is consistent with previous reports from the literature (Patton et al. 2020; Srinivasan and Alvarez 2007). To expand on these results using a different host plant, we conducted additional experiments with *M. persicae* using the host plant *N. benthamiana*. *M. persicae* that acquired PLRV from *N. benthamiana* tissue locally infected with PLRV using transient expression or from plants systemically infected with PLRV show an increase in nymph production compared to the uninfected plant controls (*P* = 0.002, ANOVA) (Supplementary Fig. S5). We did not observe a significant difference in *M. persicae* fecundity when aphids acquire virus from either type of PLRV-infected source tissue (local vs. systemic, *P* = 0.625, ANOVA), which allowed us to use the transient expression of the P0 mutants to quantify changes in aphid fecundity as a function of F-box activity. The P0 mutants used in this study do not induce changes in aphid fecundity (Fig. 4), including E83A, with intact silencing suppressor activity, suggesting that P0 regulation of aphid fecundity is independent of its role in silencing suppression and, further, that a functional P0 F-box domain is not required for P0 to regulate aphid fecundity.

**Figure 4.**
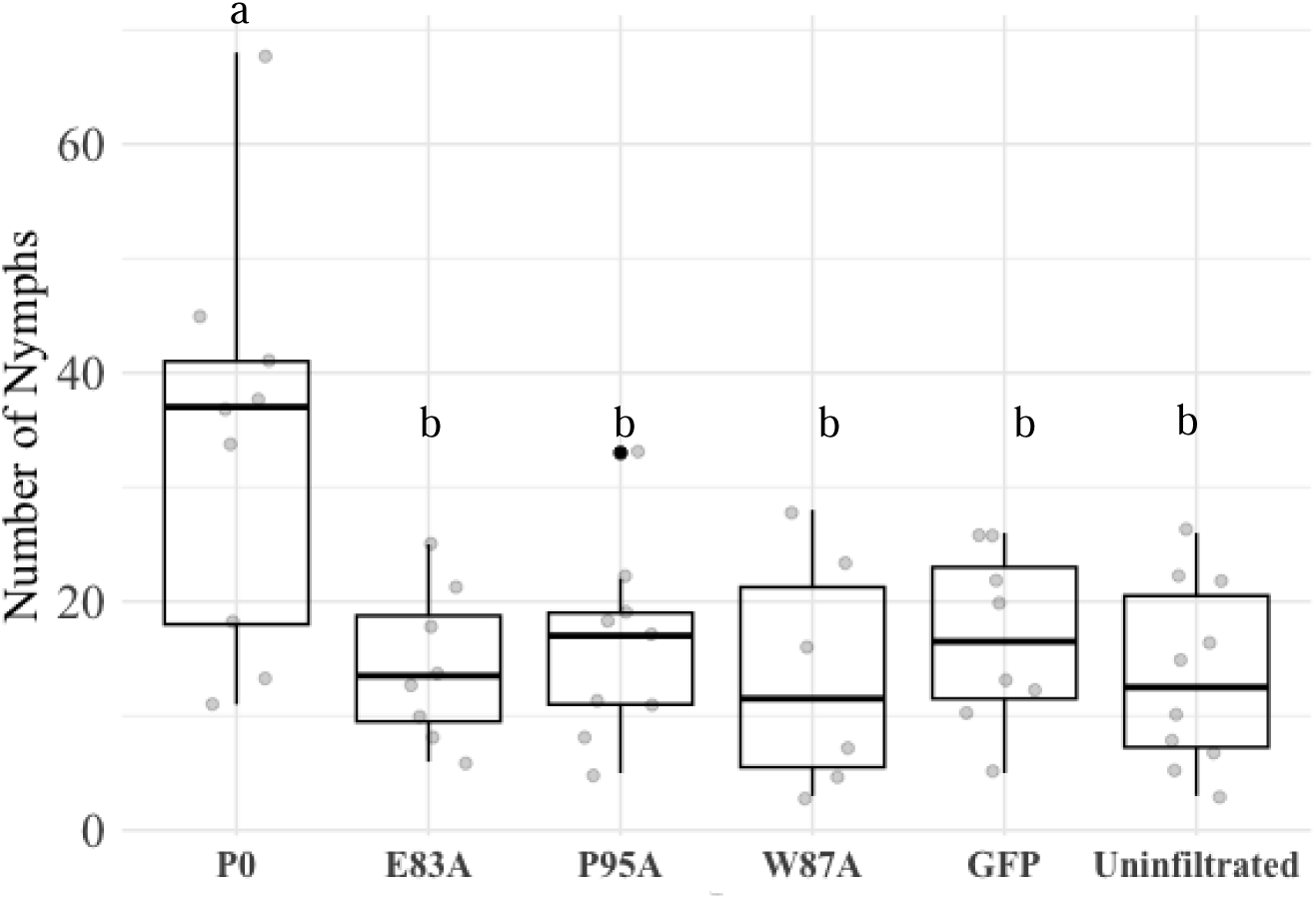
PLRV P0 wt, but not any P0 mutant, increases fecundity in *M. persicae*. To assess if P0 wt increases fecundity in an F-box dependent manner, the P0 wt and three P0 mutants (E83A, P95A, and W87A) were transiently expressed in *N. benthamiana*. Three aphids were clip caged onto *N. benthamiana* leaves for five days. The nymphs were counted, and the experiment was performed four times independently. The letters signify statistical difference via ANOVA and Tukey’s HSD.

### P0 from cotton leafroll dwarf virus is a weak silencing suppressor and does not increase MpDNV titer in *M. persicae*

To explore the effects of a closely related polerovirus silencing suppressor protein on *M. persicae* anti-viral immunity, we tested the P0 protein from the cotton leafroll dwarf virus (CLRDV) Texas isolate, accession MN872302.2. The P0 proteins of CLRDV and PLRV aligned share approximately 22% conserved residues, six of those in the F-box motif, including the mutated residues in PLRV P0 L76 and P95 (Fig. 1). The studied isolates of CLRDV P0 are classified as weak silencing suppressors (Akinyuwa and Kang 2024; Akinyuwa et al. 2023). In our observations using confocal microscopy (Table 1), fluorescence was observed, though there is no significant difference in fluorescence intensity of CLRDV P0:GFP:dsGFP compared to GFP:dsGFP (average fluorescence intensity = 6.493 RFU, *P* = 0.2310), confirming its weak silencing suppression activity. Transient expression of CLRDV P0 does not increase the titer of MpDNV in *M. persicae* compared to the un-infiltrated control and the GFP control (Kruskal-Wallis test, *P* = 0.8266, Fig. 5).

**Figure 5.**
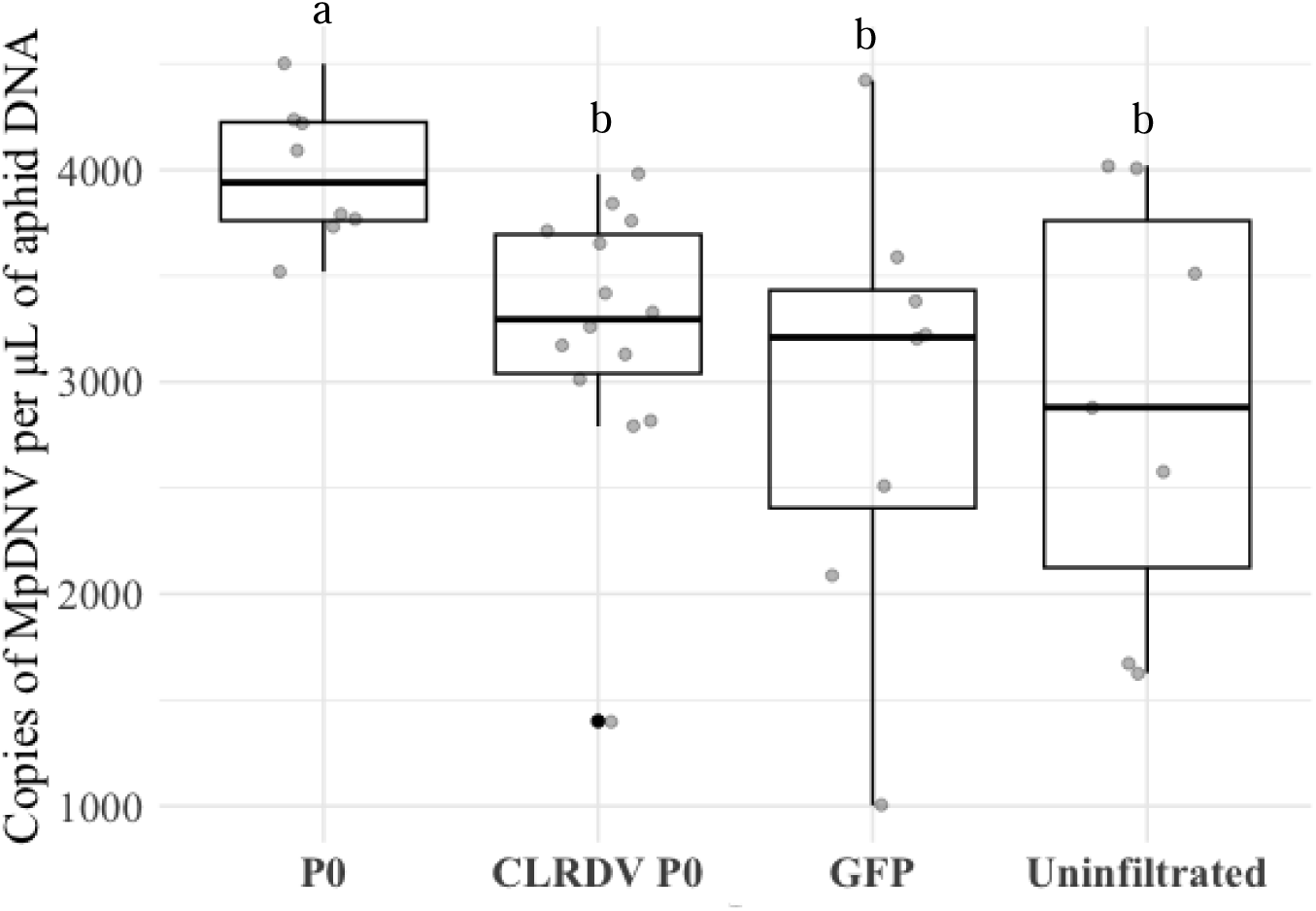
The effect of a weak and strong polerovirus P0 protein on *Myzus persicae* densovirus (MpDNV) titers. The MpDNV titers of aphids fed on *N. benthamiana* plants transiently expressing PLRV-P0, CLRDV-P0, GFP, and an un-infiltrated control for two days. Aphid DNA was extracted and MpDNV titer was quantified via droplet digital PCR, three pooled aphids in each replicate, at least ten replicates per treatment. Letters indicate significantly different treatments via ANOVA and Tukey’s HSD.

### Myzus persicae densovirus (MpDNV) is inoculated into *N. benthamiana* tissues after aphid feeding

MpDNV virions have been reported in plants where *M. persicae* had previously fed and colonized (Van Munster et al. 2003). We hypothesized that specific effects of P0 on MpDNV titer in aphids may alter MpDNV stability/retention in the plant following aphid feeding. To test this hypothesis, a PCR analysis was conducted and shows detectable MpDNV DNA in low-density, aphid-exposed plant tissues (three aphids per plant) transiently expressing P0 and the E83A mutant (which did not abolish silencing suppressor activity, Supplemental Fig. S6a). MpDNV DNA is not detected in aphid-exposed plant tissue expressing the full infectious clone of PLRV, the P95A P0 mutant that abolished silencing suppressor activity, or in control plant tissue (Supplemental Fig. S6a). When plants are overcrowded with *M. persicae* (approximately 50 aphids confined to a single clip cage), MpDNV DNA is detectable in the caged leaf areas and systemically and in newly developing leaves of all treatment and control plants, including PLRV, P0 wt, P95, and un-infiltrated plants. (Supplemental Fig. S6b).

### Other VSRs modulate aphid fecundity and founder aphid survival but not MpDNV titer

The changes in MpDNV titer and aphid fecundity due to P0 wt expression led us to test whether these interactions were specific to modulation by the VSRs of aphid-borne, circulative plant viruses or whether aphids are generally sensitive to any perturbation in the plant host RNAi pathway. We examined MpDNV titer and aphid fecundity after feeding aphids on plant tissue expressing a panel of well-characterized silencing suppressor proteins: tomato bushy stunt virus P19 (P19) (not aphid-borne), turnip crinkle virus p38 (p38) (not aphid-borne) and cucumber mosaic virus 2b (2b) (aphid stylet-borne) that target other components of the plant’s RNAi machinery (Iki et al. 2017). Although in planta, transient expression of all tested silencing suppressors increase the MpDNV titer in aphids, only aphids exposed to plant tissue transiently expressing PLRV P0 wt have significantly higher MpDNV titers as compared to aphids feeding on the un-infiltrated control (*P* = 0.0363, Fig. 6). The P19 and 2b silencing suppressor proteins increase MpDNV but with more variability than P0 wt compared the un-infiltrated control (*P*-value = 0.079). The p38 silencing suppressor does not significantly increase the MpDNV titer in *M. persicae* compared to the un-infiltrated control (*P*-value = 0.1069, Fig 6). In assays measuring fecundity rates, transient expression if the 2b protein significantly increases aphid fecundity over time. The greatest total number of aphid progeny is observed on leaves transiently expressing 2b, followed by PLRV P0 (Fig. 7a). The 2b treatment is the only treatment that was significantly different than the un-infiltrated control in the long-term fecundity assays (*P*-value = 0.0372, Fig. 7a).

**Figure 6.**
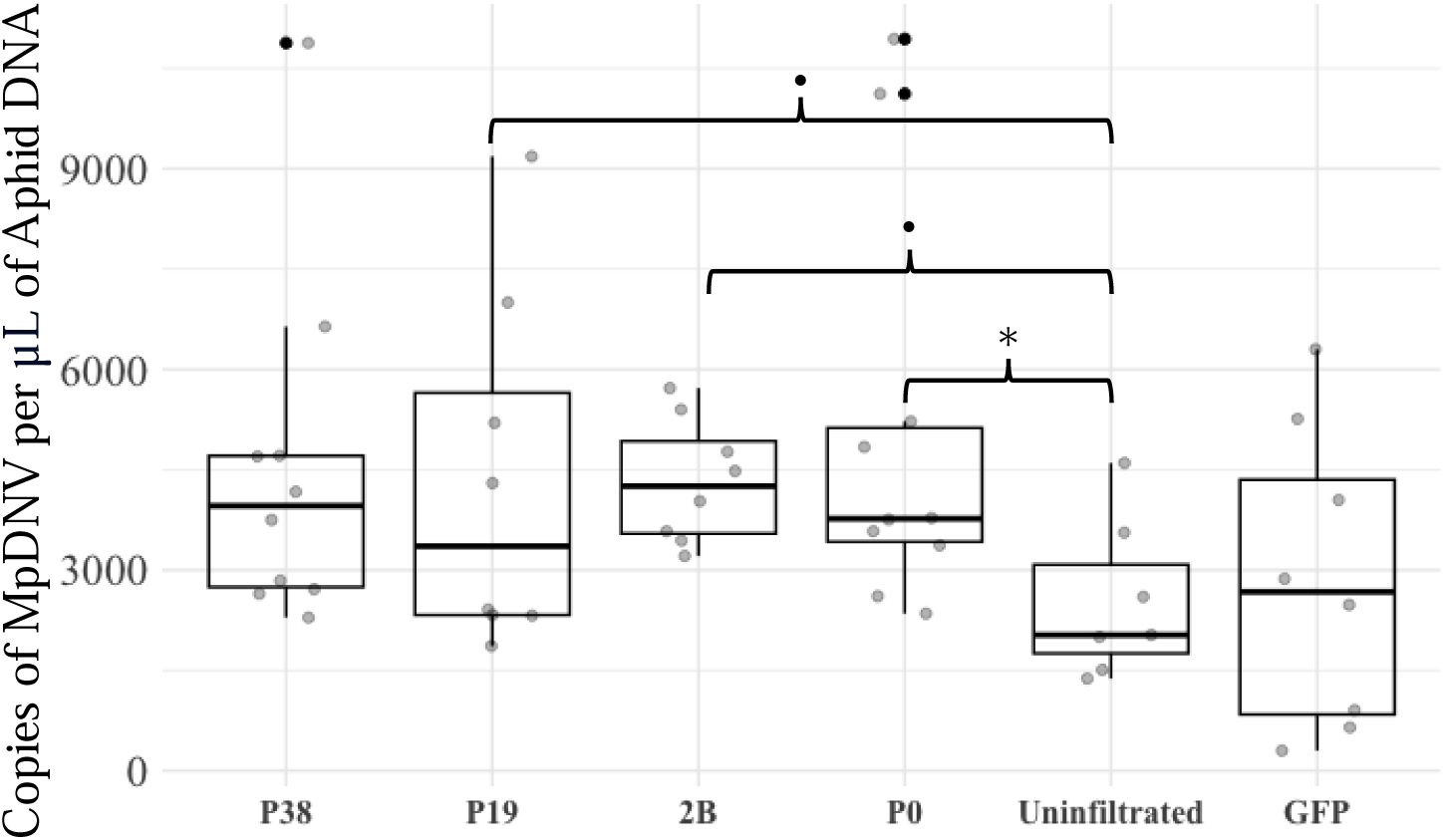
The effect of a panel of viral silencing suppressors targeting the smallRNA pathway in plants, on *M. persicae* MpDNV titers. Aphids were synchronized and fed on *N. benthamiana* leaves transiently expressing four silencing suppressor constructs; tomato bushy stunt virus P19, PLRV P0, turnip crinkle virus p38 (TCV p38), cucumber mosaic virus 2b (CMV 2b), GFP, and an un-infiltrated control for three days. Extracted aphid DNA was quantified for MpDNV titers via droplet digital PCR, three pooled aphids in each replicate, at least ten replicates per treatment. A linear mixed model was used to understand the effect of the silencing suppressor proteins on MpDNV titer in aphids, the dots represent *P*-values less than 0.1 and greater than 0.05, and the asterisk represent *P*-values less than 0.05.

**Figure 7.**
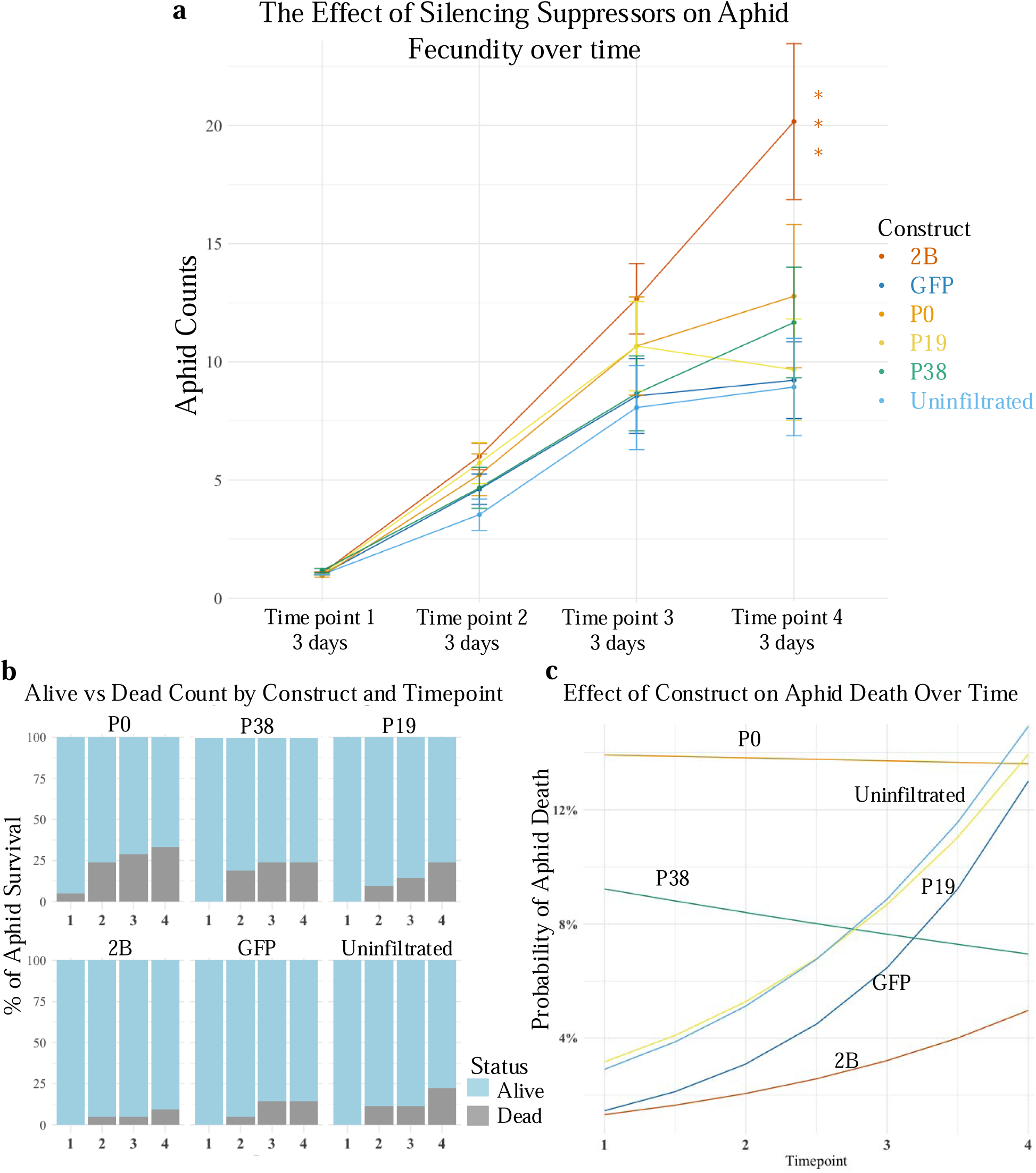
Overtime aphid fecundity increases as aphids feed on *N. benthamiana* tissue expressing VSRs. To assess if VSRs from viruses transmitted in different modes affect fecundity, *N. benthamiana* plants were Agro-infiltrated with TBSV P19, PLRV P0 wt, TCV P38, CMV 2b, and GFP. One aphid (founder aphid) per infiltration zone was independently clip caged and the total nymphs were counted every three days. **A)** Aphid fecundity measured as Aphid Counts over the four timepoints (3, 6, 9, and 12 days). The data was analyzed using a linear mixed model in the package lmer in R. **B)** The founder aphid’s survival was documented at each timepoint and treatment across the fecundity experiment (n=21, per treatment). **C)** The effect of each construct on the probability of aphid death over time. To predict the probability of aphid death per construct overtime, the survival analysis data were fit using the R package glm.

To examine the founder aphid survival as function of the expression of each VSR, we measured aphid survival from the different trials (Fig. 7b). A founder aphid is the first age-synchronized aphid at the start of each experiment. To analyze if the treatments affect death over time, we calculated and plotted the predicted probability of death for each treatment at each timepoint (Fig. 6c). The predicted probability of death for the controls (un-infiltrated and GFP) followed a consistent uptrend, suggesting the probability of aphid death increases steadily overtime as the insect naturally ages. P0 wt has a higher probability of death on the onset of the experiment, with an even, linear trend, suggesting that regardless of time, feeding on P0 wt increases the mortality of the founder aphid. Interestingly, p38 shows a slight downward trend, with a lower probability of aphid death. Finally, 2b has the lowest probability of aphid founder death regardless of the timepoint, consistent with previous findings showing that 2b decreases aphid mortality when expressed *in planta* (Ziebell et al. 2011).

## Discussion

Previous work established that PLRV P0 (P0 wt as referred to in our study), a VSR, alters aphid anti-viral immunity to an insect-infecting virus, MpDNV (Pinheiro et al. 2019). Aphids exposed to PLRV-infected plants generate unusually long sRNAs and a reduced number of 22-mers against MpDNV. These aphids have a higher titer of MpDNV than aphids exposed to un-infected plants (Pinheiro et al. 2019). In this study, we built on that prior work to investigate the impact of mutations in the P0 F-box domain, along with a panel of well-studied VSRs on MpDNV titer in aphids, aphid fecundity, and aphid survival. Collectively, the results indicate that these aphid phenotypes are responsive to VSR expression in plants.

We characterized the effect of single amino acid mutations of the PLRV P0 F-box motif on local RNA silencing in *N. benthamiana*. Prior to this study, only a few mutated residues (L76, W87, G88) were known to abolish suppressor activity (Bortolamiol et al. 2007; Zhuo et al. 2014). We generated five PLRV P0 F-box mutants: L76R, L76A, P95A, E83A, and W87A and used transient expression assays and fluorescence microscopy to quantify their silencing suppressor activity. Consistent with the findings reported with the Argentina PLRV P0 strain (Barrios Barón et al. 2021), the L76R and W87A mutants from the P0 wt sequence of our infectious PLRV clone significantly decreased the activity of P0 (Fig. 2). L76A and P95A dampened the silencing suppressor activity, while E83A had no effect on silencing activity (Fig. 2). The AlphaFold structure of P0 wt (Supplementary Fig. S1) predicts that the altered amino acids in impaired mutants are embedded within the structure, whereas the altered residue in the E83A functional mutant is surface-exposed. The buried residues of the F-box motif may be critical for the structural integrity of P0 and RNA silencing capabilities. Interestingly, a study conducted with nightshade plants, natural hosts of PLRV, showed that PLRV infectious clones carrying the P0 L76F and W87R mutations had varied accumulation patterns in plants. The PLRV-L76F strain accumulated in the inoculated leaves whereas the PLRV-W87R did not (Rashid et al. 2019). Thus, some F-box mutations may not completely disrupt the structure but rather impact virion and/or viral protein accumulation within plant cells.

The surface exposed E83A mutant, with functional suppressor activity, increased MpDNV titer in aphids and the range of densovirus titer values compared to non-functional mutants. The P95A and W87A mutants, with residues predicted to be buried within the folded P0 protein, abolished the effects of P0 on MpDNV titer in aphids. Our results support a dynamic model where P0 silencing suppressor activity in plants, regulated by the F-box motif, influences antiviral immune responses to MpDNV within aphids. We hypothesize that P0 is impacting the aphid indirectly through the plant, via the downstream effects of the silencing suppressor protein expression. Together, these data show two distinct functions for PLRV P0 in mediating interactions with its vector, *M. persicae*, one that is dependent on F-box mediated silencing suppression activity and one that is independent of that activity. To date, we cannot determine if P0 is also ingested by aphids and exerts direct effects in aphid tissues independently from its role as a VSR in plants from these experiments.

Infectious MpDNV virions have previously been reported in plant tissue (Van Munster et al. 2003). Our study builds on these observations with data supporting the hypothesis that P0 expression and aphid density are key components for MpDNV transfer from aphids into plant tissues. We detected MpDNV in plant DNA extracts when P0 wt or the functional E83A mutant were expressed, but minimal MpDNV was detected in plant tissues expressing the WT PLRV infectious clone or the P0 null mutant P95A. At low aphid densities, expression of P0 wt resulted in higher levels of MpDNV in plant tissue at the aphid feeding site. At higher aphid densities, MpDNV was found systemically in the plant, independent of P0 wt. These findings suggest that the role of P0 in signaling to aphids for antiviral immune suppression may be most functionally and ecologically significant during the initial encounter between aphids and PLRV-infected plants when aphid density is low. During PLRV infection, a scenario may involve P0 wt expression mediating changes in MpDNV titer in aphids and plants and subsequent aphid population increases, which leads to an increase in the dispersal of both viruses. Higher titer of MpDNV in aphids, whether through plant ingestion or via suppression of aphid anti-viral immunity would signal the formation of winged morphs (an aphid behavior already correlated with overcrowding), an increase in aphid reproductive rate, aphid dispersal and hence greater plant-to-plant virus transmission. Our results and this hypothesis are consistent with previous observations on aphid-densovirus interaction (Rozo Lopez and Parker 2023; Ryabov et al. 2009).

Manipulation of aphid vector characteristics by PLRV involves changes in plant physiology and the actions of several viral proteins. The polyphenic nature of aphids allows for the long-range spread of PLRV, with winged aphids being able to colonize many plant species in fields. Studies show non-viruliferous aphids preferentially settle on virus-infected plants due to virus-induced changes in the plant (Eigenbrode et al. 2002). Aphid fecundity rates significantly increase when placed on PLRV-infected plants (Castle and Berger 1993). An increase in fecundity coupled with preferential setting optimizes PLRV spread across a habitat or field. The PLRV P0, P1, and P7 proteins independently increase aphid development rate (Patton et al. 2020). These proteins mediate changes in plant-aphid interactions such as the inhibition of aphid-induced production of jasmonic acid in plants (Patton et al. 2020). The specific role of the PLRV P0 F-box motif, if any, on the inhibition of aphid-induced salicylic acid, jasmonic acid, and ethylene production in plants has not been tested, but is an important component of future work.

To measure the impact of another VSR from a polerovirus, we tested the effect of the wild type P0 protein from the cotton leafroll dwarf virus (CLRDV P0 wt) on the impact of MpDNV titers in *M. persicae*. CLRDV is not transmitted by *M. persicae*, although it is transmitted by *Aphis gossypii* (cotton aphid) in a similar, circulative, non-propagative mode. Our results indicate that CLRDV P0 wt does not increase the MpDNV titer in *M. persicae*, compared to PLRV P0 wt or the controls. CLRDV P0 wt has previously been classified as a weak silencing suppressor protein (Agrofoglio et al. 2019) and is unable to inhibit the spread of systemic silencing signals in plants (Delfosse et al. 2014). In contrast, PLRV P0 wt is considered a strong VSR in our study and is effectively able to inhibit systemic silencing signals in plants (Rashid et al. 2019). Future studies should investigate the effects of CLRDV P0 wt on A. gossypii-specific viruses or reproductive rates to determine whether the absence of observed effects on M. persicae is due to its inability to transmit CLRDV.

It is plausible to hypothesize that the VSR potency will impact the magnitude of impacts on aphid vectors. However, potency of the VSR alone is not the sole determinant of influencing aphid physiology, but rather aphids are responding to different molecular signals in the plant’s RNAi pathway. Our data show that other strong VSRs, such as the 2b, P19 and p38 proteins have different effects on aphid fecundity, survival and MpDNV titer. The p38 protein suppresses RNA silencing by preventing RISC assembly, binding to dsRNA and inhibiting siRNA from joining RISC (Liu et al. 2023). P19 is a symptom determinant in viral infection and sequesters small interfering RNAS (siRNAs). CMV 2b is a symptom determinant and interacts and inhibits AGO1, a major component of the RISC pathway and the same host target of PLRV P0 (González et al. 2010).

The connection between the aphid and plant RNAi pathway has been extensively leveraged in aphid functional genomics studies (Christiaens and Smagghe 2014; Coleman et al. 2015; Guo et al. 2014; Mulot et al. 2016; Pitino et al. 2011; Zhang et al. 2022; Zhao et al. 2018). Expression of dsRNAs that target aphid genes in plants has been shown to be an effective means of silencing aphid genes (Pitino et al. 2011; Zhao et al. 2018). Similarly, plant viruses have been engineered to express fragments of insect genes (Kolliopoulou et al. 2020). During plant infection with plant viruses engineered to express insect (including aphid) gene fragments, aphids and other insects acquire the siRNAs produced by the plant during infection with the modified virus, which modulates gene expression and phenotypes of the insects feeding on these plants (Hajeri et al. 2014). Our work shows that in the absence of heterologous expression of aphid genes, aphid physiology is sensitive to changes in specific components of the plant RNAi pathway, in particularly aspects of the pathway regulated by AGO1. Collectively, our results suggest that direct modulation of AGO1 by VSRs, such as P0 and CMV 2b, have the most profound effects on aphid interactions as compared to the VSRs p38, that inhibits siRNA loading onto AGO proteins (Iki et al. 2017). Further research is needed to better understand the evolutionary and molecular mechanisms hijacked by plant viruses that coordinate the aphid and plant RNAi pathways to understand whether these changes enable the aphid to sense signals from the plant’s RNAi pathway during plant virus infection to regulate its own development and behavior.

## Methods

### Aphid rearing conditions

Experiments in this study used parthenogenetic colonies of the green peach aphid, *M. persicae* cv. Sultz, originally collected in New York and raised on *Nicotiana benthamiana* at 20° C with an 18-hour photoperiod. To age synchronize *M. persicae*, fourth instar aphids were placed on three-week-old *N. benthamiana* plants for 48 hours to give live birth. After 48 hours, the fourth instar aphids were removed, and the nymphs were left on the plants to develop for 5 days.

### Plasmid construction

The DNA fragments containing P0 mutants, and the panel of VSR sequences were synthesized by Twist Biosciences for Gateway cloning, with sequences incorporating AttB1 and AttB2 sequences for PCR amplification required for Gateway cloning (Xu and Qingshun 2008). After gene synthesis, the fragments were diluted to a concentration of 200 ng/μL. The diluted fragments were then individually recombined into pDONR207 vectors using BP recombination reactions facilitated by Gateway BP Clonase II enzyme mix (Thermo Fisher Scientific). Following successful cloning, all clones underwent Sanger sequencing to confirm sequence integrity. Subsequently, sequence-verified clones were recombined into the destination vector pEearlygate100 via LR recombination reactions, following the manufacturer’s protocol with Gateway LR Clonase II enzyme mix (Thermo Fisher Scientific). Transient expression of all VSRs are driven by the 35s promoter (Odell et al. 1985). The resulting pEearly100 destination vectors containing the VSRs and P0 mutants were transformed into *Agrobacterium tumefaciens* strain GV3101 via electroporation.

### Aphid DNA extraction

*M. persicae* DNA was extracted from individual and pooled aphids using the Qiagen Blood and Tissue kit with the following protocol modifications: The tubes were submerged in liquid nitrogen and then subject to cryogrinding at 27 Hz for 3 minutes in a Retsch Mixer Mill 400 and resuspended in Qiagen ATL buffer, vortexed and the supernatant was moved to a new 1.5 mL tube. A total of 10 uL of Qiagen Proteinase K was added to 1.5 mL tube and incubated at 56 °C for 40 minutes. Qiagen Buffer AL was added followed by an equal part of ethanol 190 proof. The entire mixture was loaded into the DNeasy Mini spin column and collation tube and centrifuged at 8000 rpm for one minute at room temperature. Buffer AW1 followed buffer AW2 was added. The DNA was eluted in 30 uL of buffer AE, quantified by nanodrop for digital droplet PCR (ddPCR) analysis, and stored at −20 °C immediately.

### Quantifying MpDNV titer using digital droplet PCR

The digital droplet PCR (ddPCR) assay was developed for MpDNV by (Pinheiro et al. 2019), and was slightly modified in this study for virus titer measurements on the QX200 droplet digital PCR system (Bio-Rad) as follows. The ddPCR reaction for absolute quantification of MpDNV was as followed: 10 μL of 2X ddPCR Evagreen SuperMix (Bio-Rad), 1 μL of each 10 μM MpDNV primers (5’-TGACAATGGGTATATTCATTGACCT-3’ and 5’-ATCGTGCGTCAAAAGAAACCCT-3’), 7 μL of dH_2_O and 2 μL of DNA at 1ng/uL and added to a 20 μL reaction. A cartridge holder containing 20 μL of the ddPCR reaction and 70 μL of droplet generator oil for Evagreen (Bio-Rad) was placed into the QX100 droplet generator (Bio-Rad) where 40 μL droplets were generated. Droplets were transferred to a 96-well plate (Eppendorf) and sealed with easy pierce foil (Bio-Rad). The PCR amplification was carried out on the Applied Biosystems 2720 Thermocycler. The thermocycling conditions began at 95°C for five minutes, followed by 40 cycles of 95°C for 30 seconds and 60°C for one minute, one cycle at 4°C° for five minutes, one cycle at 90°C for five minutes and ending at 12°C. Following amplification, the plate was placed into the droplet reader cassette and loaded into the droplet reader (Bio-Rad). The ddPCR droplet data were analyzed using the QuantaSoft analysis software (Bio-Rad), which displays the target results as copies per μL of PCR mixture.

### Silencing suppressor assays in *Nicotiana benthamiana*

To assess silencing suppressor activity of the P0 wt, P0 mutants, and the panel of VSRs, 3-week-old *N. benthamiana* plants were co-infiltrated with recombinant *Agrobacteria tumefaciens* cultures for transient expression of GFP, double stranded (ds)GFP, and one of the VSRs or P0 mutants in a 1:1:1 ratio. The *A. tumefaciens* cultures with each construct in a binary expression vector were diluted with infiltration buffer (10 mM MgCl_2_, 10 mM MES pH 5.5, 150 µM acetosyringone), to a final OD_600_ within 0.05+/− from every other culture, with a target OD_600_ of 0.315. Transient expression followed the plant inoculation protocol as described (Deblasio et al. 2018). The silencing suppressor activity via GFP expression was measured using a hand-held UV lamp at four DPI.

### Confocal microscopy and GFP expression quantification

The triple infiltrated leaf tissue of each VSR and P0 mutant was examined using confocal microscopy. A thin layer of epidermal cells was cut from the fully developed infiltrated leaves. The epidermal peels were viewed under a Leica TCS-SP5 (Leica MicroSystems Exton) confocal microscope. GFP was excited with the 488-nm line of a multiline argon laser with emission spectra collected by a hybrid detector (HyD). All scans were conducted sequentially with line averaging of eight. The fluorescence intensity was calculated from more than 20 cells (images) per treatment using ROI measuring RFU using ImageJ (need software manufacturer info and version). The relative intensity of each image was analyzed by ANOVA in R.

### Aphids feeding on infiltrated *N. benthamiana* tissue

To measure differences in MpDNV titer in *M. persicae*, aphids were fed on infiltrated *N. benthamiana* tissue transiently expressing each VSR, mutant construct, or GFP as a protein control. For each experiment, at least three leaves of *N. benthamiana* plants were infiltrated per construct three weeks post germination. Synchronized fourth instar aphids were placed into clip cages on the infiltrated leaves at 1 day post inoculation (DPI), with three to five aphids per cage. Each treatment had at least 10 biological replicates, and each biological replicate consisted of 3 aphids collected from the clip cages. Aphids were collected at three DPI (after two days of feeding) or four DPI (three days of feeding) from each treatment. Aphids were immediately placed in a 2 mL microcentrifuge tube with two 3.2 mm stainless steel metal beads from Next Advance and placed into a −80°C freezer before DNA extraction.

### Detection of local and systemic MpDNV in *N.benthamiana*

To detect the local MpDNV deposition into in *N. benthamiana*, three aphids in a clip cage were placed onto a *N. benthamiana* leaf transiently expressing the (1) PLRV P0 wt; (2) the infectious clone of PLRV; (3) P0 E83A (4) P0 P95A, (5) and an un-infiltrated leaf. After three days, the aphids were removed, and the leaf surfaces were cleaned of honeydew with 70% ethanol. Per treatment, a sample consisting of three plant tissue punches was collected in a 2mL microcentrifuge tube with two 3.2 mm stainless steel beads was collected for MpDNV detection *in planta* using PCR. The plant DNA was extracted using the Zymo DNA extraction kit following manufacturer’s instructions. The PCR was carried out using RedTaq PCR mix, following manufacturer’s instructions and MpDNV specific primers (5’ TGACAATGGGTATATTCATTGACCT-3’ and 5’-ATCGTGCGTCAAAAGAAACCCT-3’).

To assess the systemic movement of MpDNV, approximately 50 aphids in a clip cage were placed onto a singular *N. benthamiana* leaf transiently expressing either the (1) PLRV P0wt; (2) P0 E83A; (3) P0 P95A (4) TBSV P19, or the (5) un-infiltrated control. After three days, the aphids were removed, and three plant tissue punches were taken at the site of inoculation and infestation in the clip cage, outside of the clip cage on the same leaf, and from the newest developing leaves from each treatment and control. The plant punches were frozen, and DNA was extracted from the leaf punches using the Zymo DNA extraction kit following manufacturer’s instructions. A PCR followed using Red Taq polymerase, MpDNV primers, and the extracted plant DNA.

### PLRV and P0 fecundity assays

To test if PLRV infection in plants has an impact on aphid fecundity, four PLRV-infected and four healthy potato plants (cv. Red Maria) were selected. Four clip cages were placed on each plant and a total of four adult *M. persicae* were placed on each plant for seven days, after which the number of nymphs were recorded. To test if there is a difference in local or systemic PLRV infection on aphid fecundity, two-week-old *N. benthamiana* plants were initially infected with PLRV and allowed to achieve systemic infection over the course of one week. After one week, three-week-old *N. benthamiana* plants were infiltrated with PLRV. One day following infiltration, three aphids per clip cage were confined to the area of infiltration for local PLRV infection. For systemic PLRV infection, aphids were placed on the latest fully developed leaf. Four days post-infestation the aphids were removed, and nymphs were manually counted.

To determine whether the PLRV P0 F box motif is responsible for the increase in aphid fecundity in the context of PLRV infection, we measured aphid fecundity on leaves transiently expressing PLRV P0 wt and the PLRV P0 F-box mutants. Age-synchronized *M. persicae* were placed on *N. benthamiana* leaves transiently expressing the following constructs: PLRV P0 wt, W87A, P95A, E83A, or GFP and confined to the zone of infiltration using clip cages. At five DPI, the synchronized aphids were removed, and the number of nymphs were counted manually. To collect time-point specific data of the panel of VSRs on aphid fecundity, three-week-old *N. benthamiana* plants were inoculated with *A. tumefaciens* strain GV3101 with plasmids design to express PLRV P0, TBSV P19, CMV 2b, TVC p38, or GFP for transient expression in plants. One aphid per infiltrated leaf was confined to the zone of infiltration via clip cage at one DPI, and four timepoints of nymph data were collected every three days. Death of the founder aphid, (the aphid placed in the clip cage at 1 DPI) was recorded at each timepoint of the experiment.

### Statistical analysis

For confocal microscopy image analysis, the average relative intensity was analyzed by ANOVA followed by Tukey’s Honest Significant Difference (HSD) test in R. For each experimental condition, we analyzed four biological replicates, each with at least five images each. Statistical analyses were conducted in R (v4.1.2). To assess statistical differences in average MpDNV titers among aphids fed on tissue transiently expressing P0 wt and the P0 mutants, titers were quantified using ddPCR. Due to non-normal data distribution, comparisons were made using the Kruskal-Wallis test, followed by Dunn’s post hoc test with Benjamini-Hochberg correction. To measure statistical difference among the average fecundity of aphids feeding on P0 wt and P0 mutants, and the statistical difference between CLRDV P0 wt and P0 wt, an ANOVA followed by Tukey’s HSD test was used with a cut of alpha value of 0.05. A linear mixed model in R (lme4) was used to understand the impact of a panel of VSRs on MpDNV titers in aphids (Bates et al. 2015). The model included ‘treatment’ as a fixed effect and ‘trial’ as a random effect to account for independent replicates. To measure if aphid fecundity increases when aphids are fed on plant tissue transiently expressing a panel of VSRs, a linear mixed model using the R package lme4 was used with the random effects; groups, construct:plant:cage, and construct:plant. To measure the probability of founder aphid death from each VSR treatment overtime a generalized linear regression model was used and the predicted probability of death overtime was visualized using ggplot2 based on model fits in Rstudio (Ginestet 2011).

## Acknowledgements

This work was supported by the U.S. Department of Agriculture (USDA) Agricultural Research Service project # 8062-22410-007-000-D and USDA National Institute of Food and Agriculture projects 2019-67011-29610, 2020-67013-31917 and 2023-67011-40490. We thank Chad Thomas, Lisa Scanlon, and Julie Bojanowski (Cornell University) for care of the plants and insects used in the research. We thank Lynn Marie Johnson (Cornell University, Statistical Counseling Unit) for their help and guidance on the statistical analysis of the datasets.

