## Supplemental data for "Viral silencing suppressor activity in plants modifies aphid antiviral immunity and fecundity"

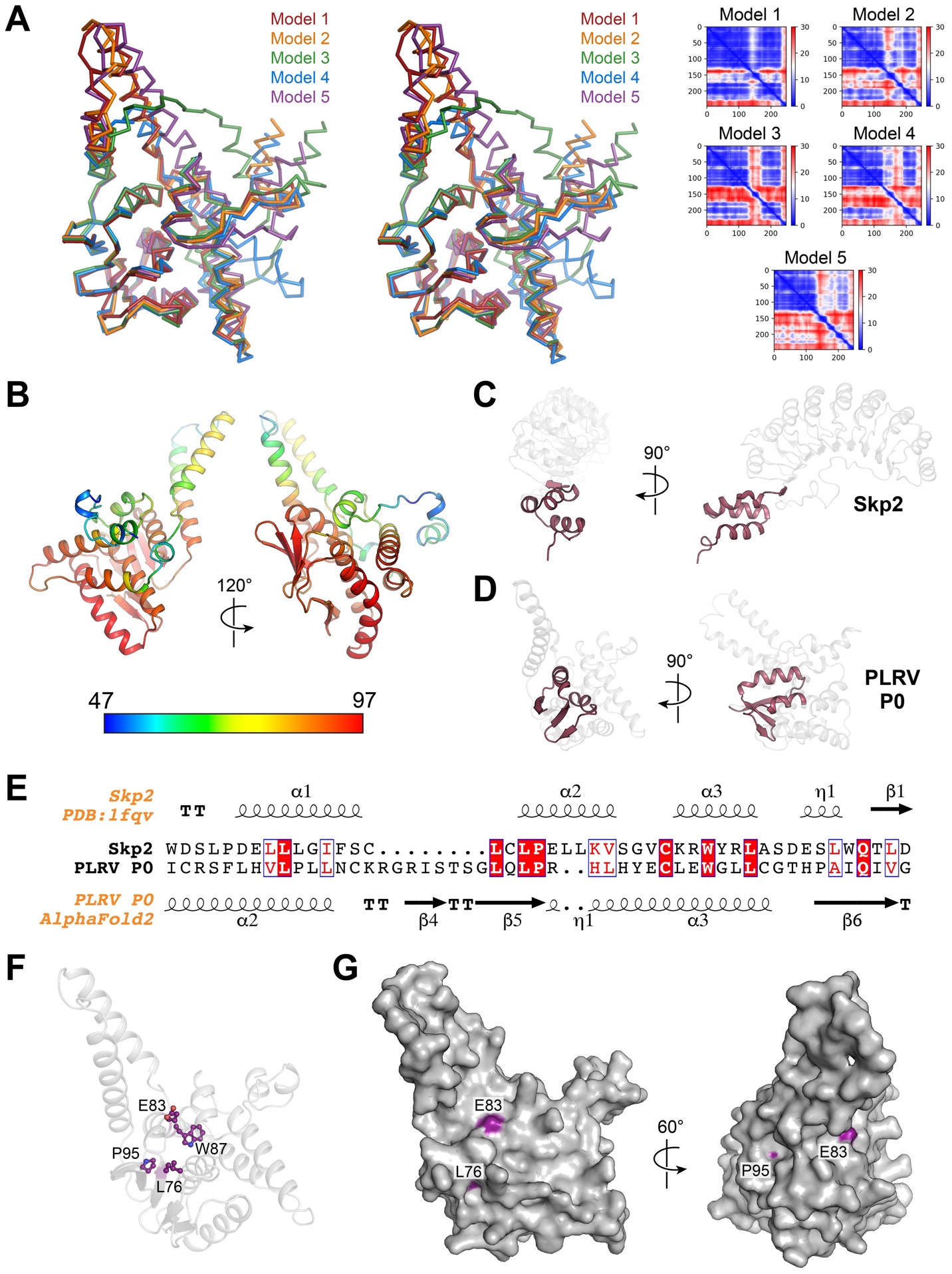

**Figure S1. AlphaFold modelling of PLRV P0.** **A**) Superposition of the top five PLRV P0 models generated by AlphaFold2 viewed in stereo(Jumper et al. 2021). Predicted aligned error plots are shown on the right. **B)** Top ranked PLRV P0 model colored according to the predicted local distance difference test (pLDDT) score (0-100), with values greater than 90 indicating high confidence and values below 50 indicating low confidence. Scale bar denotes per residue confidence coloring for pLDDT scores ranging from 47 to 97. **C,D)** Structural organization of the F-box consensus sequence (ruby) in the human Skp2 protein (PDB: 1FQV) **C)** and the PLRV P0 AlphaFold2 model **D)**. **E)** Alignment of the F-box consensus sequence from human Skp2 and PLRV P0 with associated secondary structure features mapped above and below, respectively. Shading indicates sequence conservation: white text on red background, 100% conserved; boxed red text on white background, 70% conserved. **F,G)** Location of mutated F-box residues (purple) on the AlphaFold-modeled PLRV P0 structure (gray) rendered as a cartoon **F)** and surface **G)**.

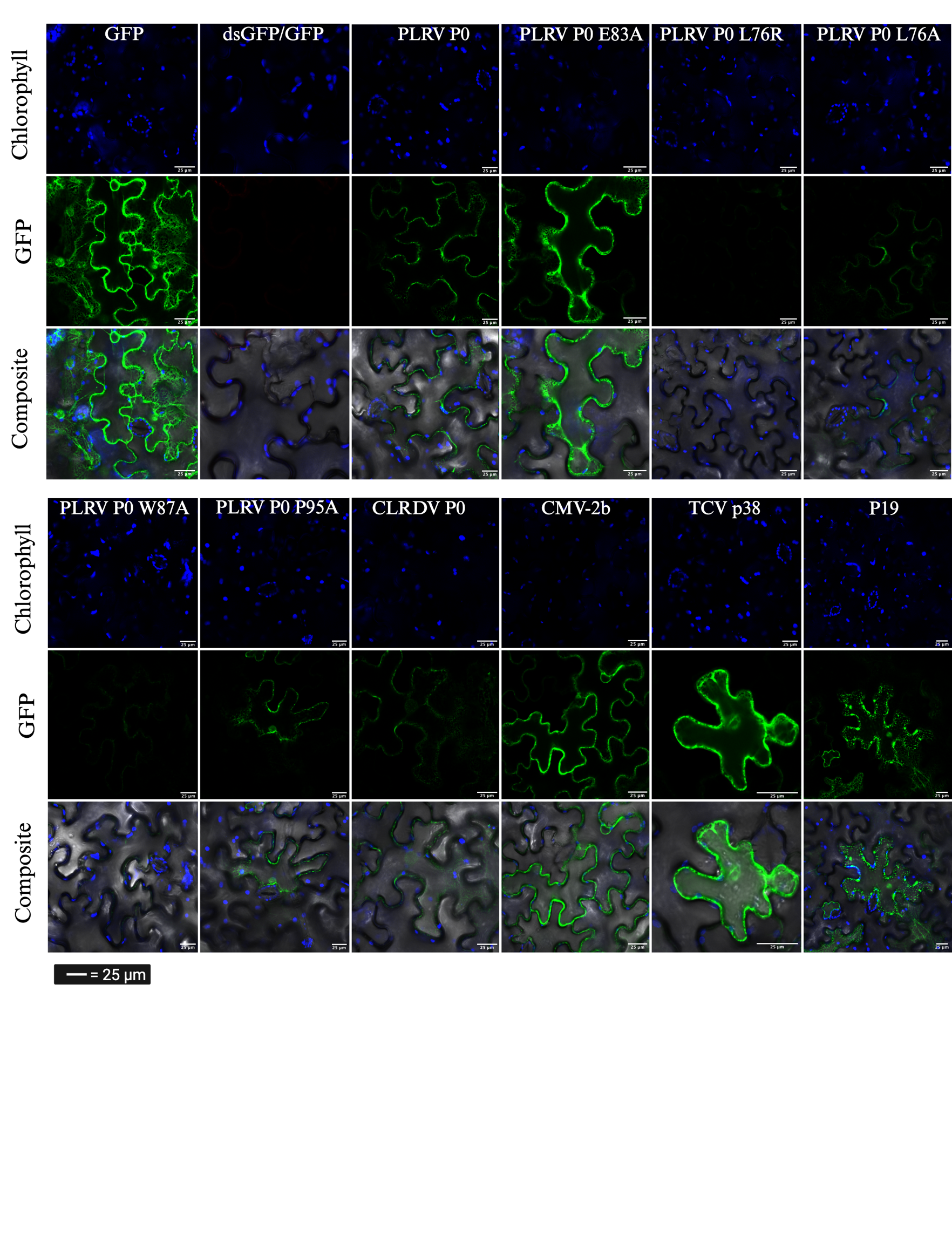

**Figure S2. Visualization of silencing suppressor activity in plants using confocal microscopy.** Confocal images of the abaxial side of *N. benthamiana* leaves transiently expressing PLRV P0 wt, PLRV P0 mutants, TCV 38p, CLRDV P0, TBSV P19, or CMV 2b, with GFP and dsGFP, in a 1:1:1 ratio to evaluate the suppressor activity of each protein. For controls, we evaluated the transient expression of GFP or GFP and dsGFP in a 1:1 ratio in *N. benthamiana* leaves. From left to right, the panels for each silencing suppressor and control are: Autofluorescence, GFP signal, and the composite.

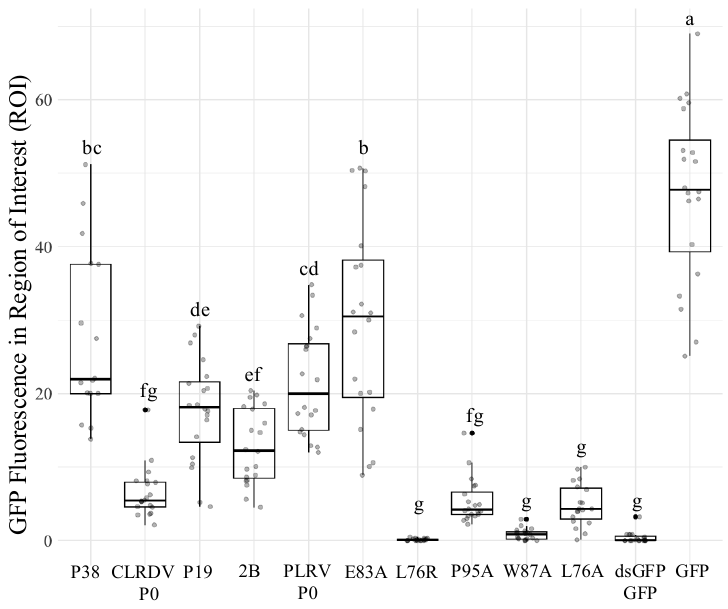

**Figure S3. The fluorescence intensity of GFP per silencing suppressor construct measured within the region of interest.**

The fluorescence intensity of GFP after the triple infiltration of GFP, dsGFP, and a silencing suppressor protein in a 1:1:1 ratio. The intensity of fluorescence signal in the infiltration zone were measured using a Leica TCS-SP5 (Leica MicroSystems Exton) confocal microscope. The fluorescence intensity of the region of interest (ROI) was calculated using the average of 20 images of cells, statistical analysis was done with ANOVA followed by Tukey’s post hoc test, letters indicate statistical significance (*P* < 0.05).

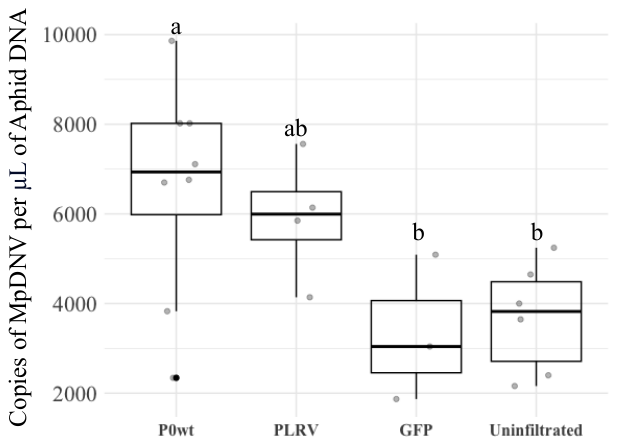

**Figure S4. P0, encoded by PLRV, increases the MpDNV titer in *M. persicae*.** Aphids fed on *N. benthamiana* plants transiently expressing the PLRV P0 wt silencing suppressor, a PLRV infectious clone, GFP, or a non-infiltrated control. Aphids were clip caged on each treatment for three days. Letters show significantly different treatments (*P*<0.05) via the Tukey’s HSD test.

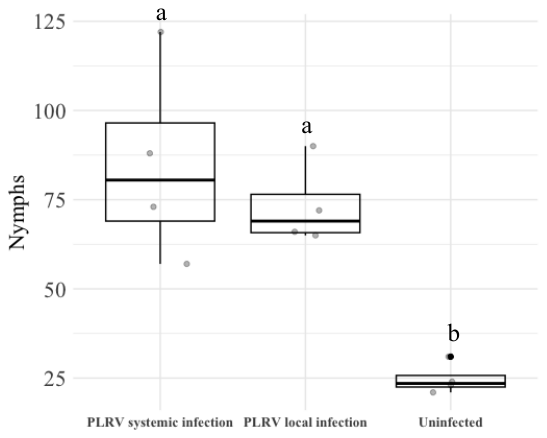

**Figure S5. Systemic and local infection of PLRV increases aphid fecundity.** To assess if the local and systemic transient expression of the PLRV-infectious clone in *N. benthamiana* increases fecundity, three aphids were clip caged onto leaves for four days, and then nymphs were counted. Local infection refers to the site of infiltration of the PLRV infectious clone, systemic infection refers to newly developing leaves infected with PLRV. The experiment was repeated three times independently, letters signify statistical difference.

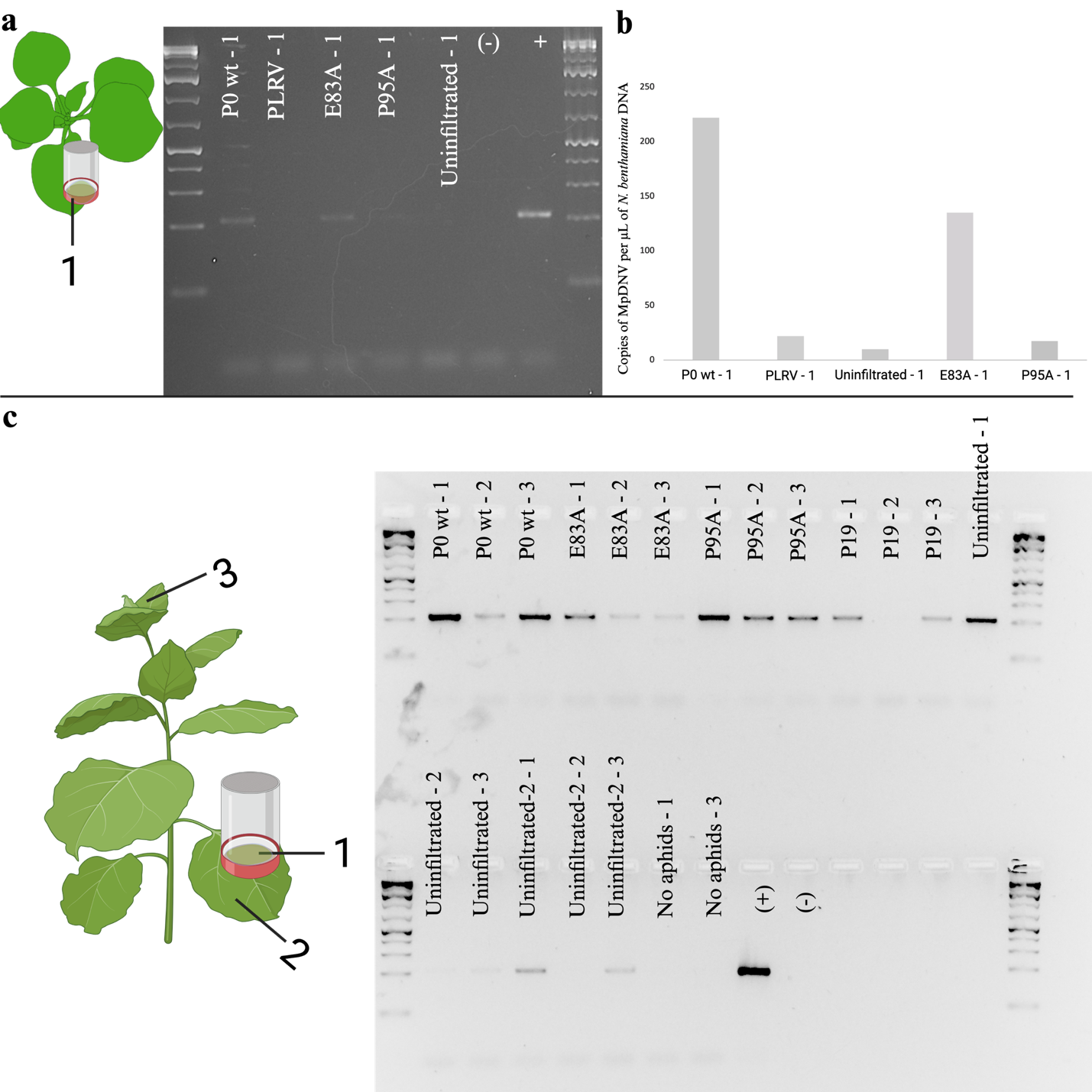

**Figure S6. Detection of Myzus persicae densovirus (MpDNV) in *Nicotiana benthamiana*. A)** Three aphids in a clip cage were placed onto *N. benthamiana* leaves transiently expressing PLRV P0wt; the infectious clone of PLRV; P0 E83A, P0 P95A, or onto an un-infiltrated leaf. After three days, the aphids were removed, and plant samples were collected for MpDNV detection *in planta* via PCR. **B**) For an absolute quantification of MpDNV in the plant samples, digital droplet PCR was run. **C)** To test for systemic movement of MpDNV in plants, approximately 50 aphids in a clip cage were placed onto *N. benthamiana* leaves transiently expressing the PLRV P0wt, P0 E83A, P0 P95A, TBSV P19, or and an un-infiltrated leaf. A leaf with a clip cage without aphids was used as a contamination control. After three days, the aphids were removed, and three plant punches were taken at the site of inoculation and infestation in the clip cage, outside of the clip cage on the same leaf, and from the newest developing leaves from each treatment and control. A PCR followed to detect MpDNV.
